# Trunk neural crest gene MOXD1 affects embryonic development

**DOI:** 10.1101/2023.08.29.555314

**Authors:** Elina Fredlund, Stina Andersson, Manon Ferreira, Sofie Mohlin

## Abstract

Ectoderm-derived neural crest is a transient structure arising during early embryogenesis in vertebrates. Neural crest consists of four derivatives based on their anterior- to posterior location along the body axis; cranial, vagal, trunk and sacral, respectively. We recently showed that trunk neural crest-specific gene *MOXD1* functions as a tumor suppressor in trunk neural crest-derived childhood cancer form neuroblastoma and is essential for proper development of healthy adrenal glands. However, the role of MOXD1 during early embryogenesis is not known. Here, we conditionally knocked out *MOXD1* in trunk neural crest cells before they become lineage-committed, using a CRISPR/Cas9 approach in chick embryos. Assessment of embryo growth showed that knockout of *MOXD1* delayed development with knockout embryos being smaller. RNA sequencing of trunk-derived neural crest cells from control and knockout embryos showed enrichment of genes connected to gland development, copper ion metabolism and neuroblastoma progression. In conclusion, MOXD1 is important during early and prolonged embryonic development with effects on gland formation, possibly mediated via its role in copper metabolism.

## Introduction

The neural crest is a transient vertebrate-specific embryonic tissue with the capacity to give rise to a plethora of cell types. The ectoderm-derived neural crest constitutes the dorsal neural tube, where pre-migratory cells form during neurulation. Upon delamination from the neural tube, the neural crest cells undergo an epithelial-to-mesenchymal transition (EMT) and start to migrate to all parts of the developing embryonic body. The neural crest is organized into four derivatives defined by the position on the axial level along the anterior-to-posterior axis; cranial, vagal, trunk and sacral. Cells originating from the trunk axial level generate for example Schwann cells, sympathetic ganglia, and chromaffin cells of the adrenal medulla (Bronner-Fraser and Fraser, 1988; Vega-Lopez et al., 2018).

Monooxygenase DBH-like 1 (MOXD1) belongs to the monooxygenase family and shares sequence homology to dopamine beta hydroxylase (DBH), an oxidoreductase catalyzing the conversion of dopamine to norepinephrine in the adrenal gland. Norepinephrine is the main neurotransmitter of the sympathetic nervous system (SNS), a structure including the adrenal gland, that is derived from the neural crest. MOXD1 is dependent on copper ions to exert hydroxyl group incorporation into its substrates. However, in contrast to the well-established function of DBH in neural crest-derived chromaffin cells, the functions of MOXD1 are still mainly unknown. *MOXD1* has been described to be enriched in the trunk neural crest of developing chick embryos as demonstrated by *in situ* hybridization and bulk RNA sequencing of sorted neural crest cells during avian development (Knecht and Bronner-Fraser, 2001; Murko et al., 2018). We recently confirmed that MOXD1 is expressed in migrating trunk neural crest also at protein level, and in addition to chick, also in human embryos (Fredlund et al., 2023). We further showed that *MOXD1* is a tumor-suppressor gene in trunk neural crest-derived childhood cancer form neuroblastoma, and that *MOXD1* affects genes connected to embryonic development in these cancer cells. Conditional knockout of *MOXD1* in trunk neural crest cells disrupted adrenal gland formation and resulted in high expression of neuroblastoma-associated protein CD56/NCAM1 at time points when all organs are fully developed (Fredlund et al., 2023). However, the effects that MOXD1 might have on early trunk neural crest- and embryonic development have not been established.

Here, we show that *MOXD1* conditional knockout in trunk neural crest cells specifically, delays embryonic development. RNA sequencing of trunk neural crest cells from embryos with or without *MOXD1* identified genes connected to cell fate commitment and gland development. Two of the most differentially expressed genes were *APOA4* and *DRR1*, both involved in copper metabolism. We conclude that MOXD1 is an important regulator of early trunk neural crest development, with a potential impact on tissue homeostasis as organs of the SNS form.

## Results

### MOXD1 knockout impedes development of embryos

Knowing that MOXD1 is expressed in trunk neural crest cells (Fredlund et al., 2023; Knecht and Bronner-Fraser, 2001; Murko et al., 2018), we set out to investigate its biological role during early neural crest development. To this end, we electroporated a morpholino or three separate CRISPR/Cas9 gRNAs for loss-of-function experiments (**Fig. 1**). Constructs were injected into the lumen of the neural tube in chick embryos at pre-migratory stages (HH10+/HH11) *in ovo*, to target trunk neural crest cells specifically and before they delaminate and go through EMT. We quantified developmental stage by two different means, HH stage by head- and tail morphology *in ovo*, or by quantifying the number of formed somites *ex ovo*. Embryos were staged 24- or 36-hours post-injection for morpholino and CRISPR/Cas9, respectively. We observed that MOXD1 knockout substantially delayed development (average HH stage 16 *vs*. 18 for morpholino, HH stage 17 *vs*. 19 for CRISPR, and 30 vs. 37 somites for CRISPR; **Fig. 2A-C)**

**Fig 1.**
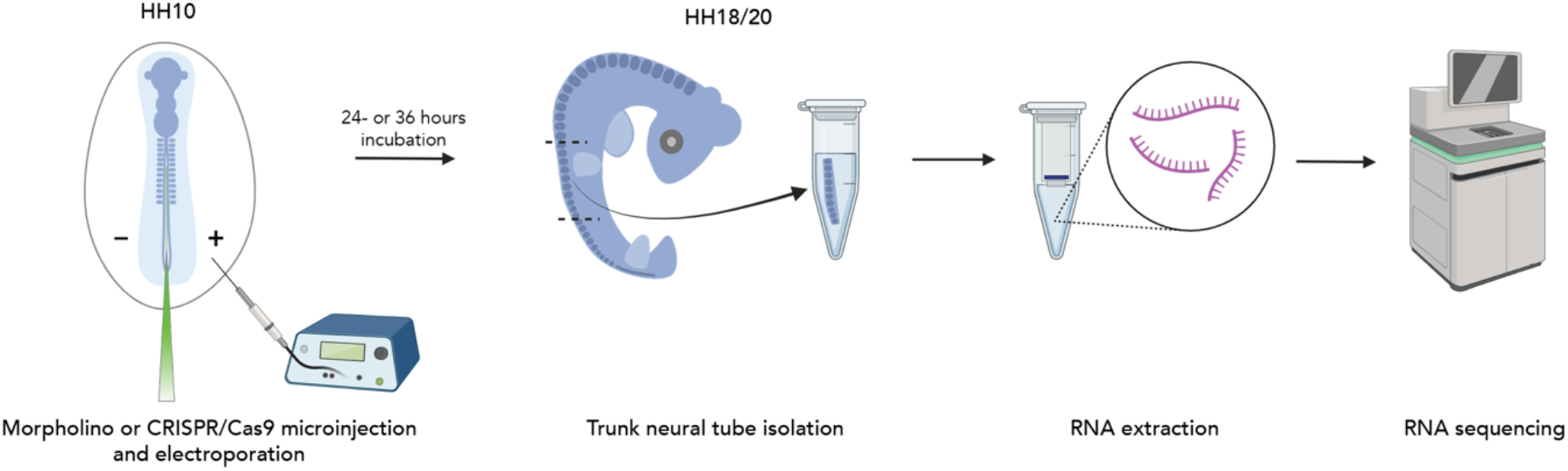
Schematic illustration of experimental set up for RNA sequencing of MOXD1 KO chick embryo. *In ovo* injection of morpholino or CRISPR/Cas9 into the lumen of the neural tube of a chick embryo at stage HH10, followed by electroporation. After 24- or 36 hours of incubation, embryos were harvested, and neural tubes isolated at the region of the trunk. RNA from individually isolated neural tubes was extracted before sending samples for RNA sequencing.

**Fig 2.**
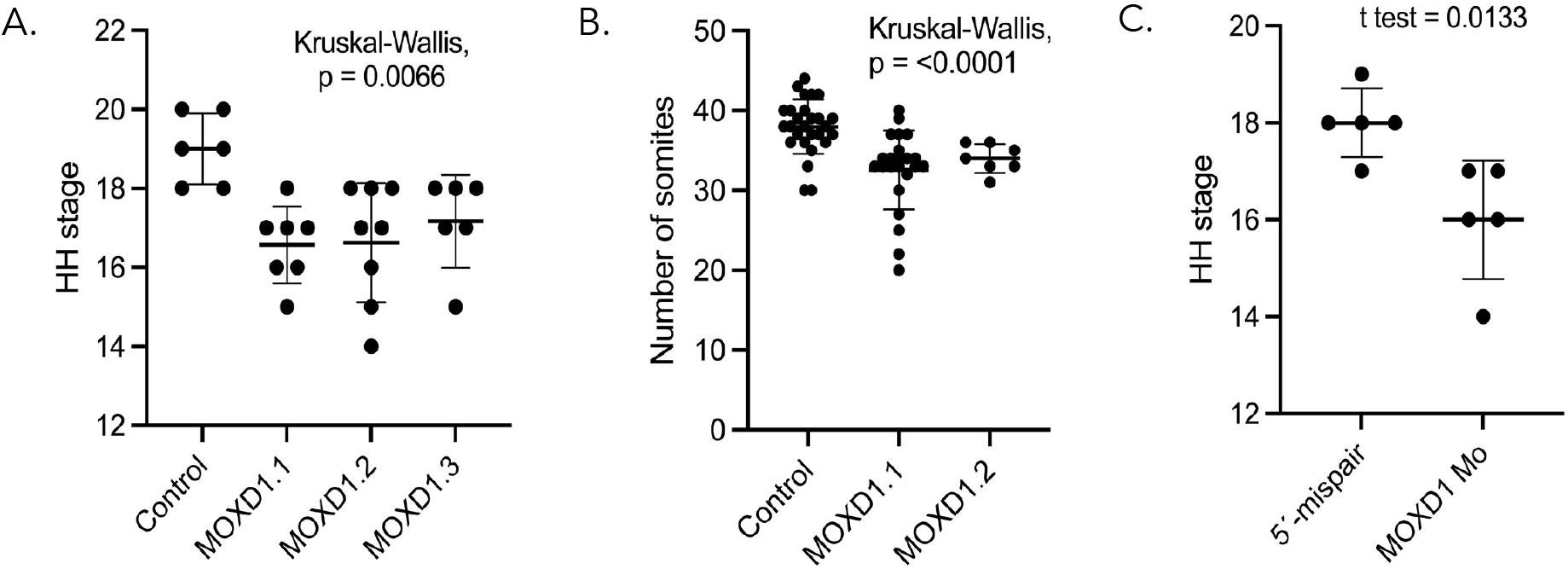
Embryo development is delayed upon loss of MOXD1 expression in trunk neural crest cells. **(A-B)** Developmental age 36 hours after CRISPR mediated knock out in trunk neural crest cells using three unique gRNAs targeting *MOXD1* at three different genomic locations. Age measured by (A) head and tail morphology converted into HH stage and (B) the number of developed somite pairs determined *ex ovo*. Number of embryos analyzed were (A) *n* = 6 (control), *n* = 7 (MOXD1.1), *n* = 8 (MOXD1.2), *n* = 6 (MOXD1.3) and (B) *n* = 27 (control), *n* = 23 (MOXD1.1), *n* = 7 (MOXD1.2). Statistical significance was determined by the Kruskal-Wallis test as indicated. **(C)** Developmental age of chick embryos determined by HH staging 24 hours post electroporation with 5’mispair or MOXD1 targeting morpholino (Mo). Number of embryos analyzed were *n* = 5 (5’-mispair), *n* = 5 (MOXD1 Mo). Statistical significance determined by Student’s t-test. (A-C) Results are presented as mean and error bars represent standard deviation.

### RNA sequencing identifies genes connected to copper binding and neural crest-related disease

To get insight into the processes governing MOXD1-driven development, we performed RNA sequencing on dissected trunk neural crest from control and *MOXD1* knockout embryos (**Fig. 1**). To enrich trunk neural crest cells, we dissected neural tube tissue at trunk axial level only. RNA sequencing identified a number of differentially expressed genes of which 24 were upregulated and 55 downregulated with *MOXD1* knockout (**Fig. 3A**). Volcano plots highlight significant and most differentially expressed hits (**Fig. 3B**). Two of these genes, *FAM107A* (also known as *DRR1*) and *NEUROD1*, have been shown to be involved in neuronal development, as well as cell growth of trunk neural crest-derived childhood cancer form neuroblastoma (Asano et al., 2010; Lu et al., 2015; Mu et al., 2017). Cell growth suppressor *FAM107A* (Mu et al., 2017) is downregulated when *MOXD1* is knocked out (**Fig. 3A-B**), in concordance with our recent findings of how MOXD1 acts as a tumor-suppressor in neuroblastoma (Fredlund et al., 2023).

**Fig 3.**
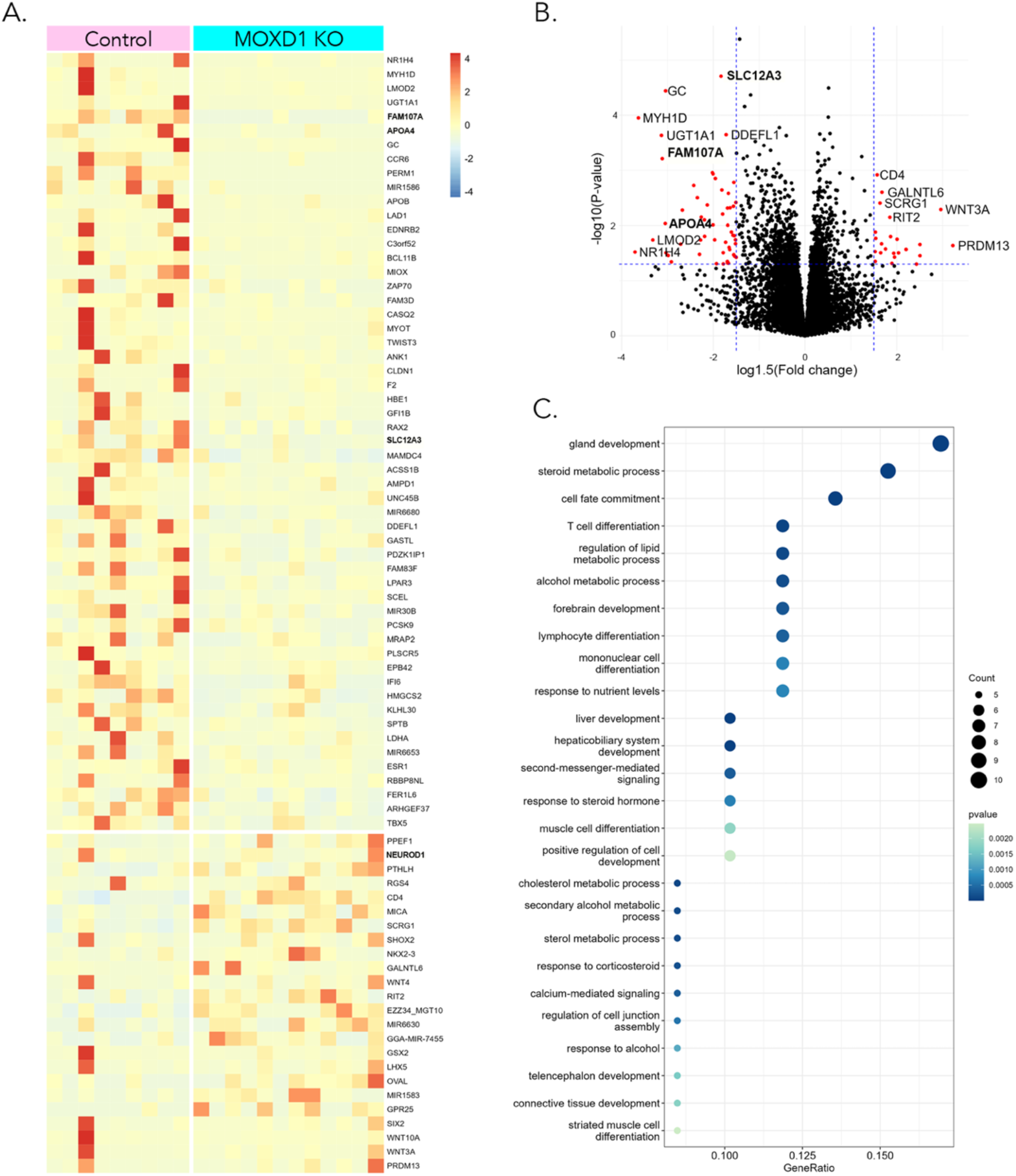
RNA sequencing reveals MOXD1 involvement in organ development and cell fate. **(A)** Heatmap of the most differentially expressed genes (DEGs) from RNA sequencing, comparing control and *MOXD1* CRISPR knockout embryos. Neural tubes from individual embryos were run as biological replicates (n=9 control and n=12 MOXD1 KO). Cut-off p <0.05 and log2Fold change >1.5. **(B)** Volcano plot of control embryos and *MOXD1* CRISPR knockout embryos. Blue lines represent cut-off p <0.05 and log2Fold change >1.5, respectively. **(C)** Gene set enrichment analysis of the DEGs from (A).

The *NEUROD1* gene is upregulated when *MOXD1* is knocked out (**Fig. 3B**), in line with MOXD1 acting as a tumor-suppressor (Fredlund et al., 2023), and NEUROD1 promoting neuroblastoma cell growth (Lu et al., 2015). Considering that MOXD1 also relates to copper function, it is notable that the copper ion-binding gene *APOA4* (Wong et al., 2007) is downregulated when *MOXD1* is knocked out (**Fig. 3B**). Of note, FAM107A has been shown to form complex with F-actin and COMMD1, where the latter gene has an important function in copper metabolism (Mu et al., 2017).

In our recent paper showing that MOXD1 is a tumor suppressor in neuroblastoma, we performed RNAseq on tumor cells grown *in vitro*, and after they had formed tumors *in vivo* (Fredlund et al., 2023). To check for putative overlaps between healthy development and cancer, we crossed our lists of DEGs from all three RNAseq experiments. We extracted five genes, *APOA4, DUSP15, IMPG2, RBBP8NL*, and *TDRD9* (**Table 1**).

**Table 1.** List of genes overlapping with RNA sequencing from Fredlund et al (2023)

| **Gene** | **Gene description** | **Human chr** |
| --- | --- | --- |
| **IMPG2** | Interphotoreceptor matrix proteoglycan 2 | 3q |
| **TDRD9** | Tudor domain containing 9 | 14q |
| **RBBP8NL** | RBBP8 N-terminal like | 20q |
| **APOA4** | Apolipoprotein A4 | 11q |
| **DUSP15** | Dual specificity phosphatase 15 | 20q |

### MOXD1-regulated genes are involved in organ development

To investigate which cellular processes MOXD1 affects during development, we performed gene set enrichment analysis. Several pathways were shown to regulate organ development, including that of glands (**Fig. 3C**). In addition, *MOXD1* knockout enriched genes involved in cell development and cell fate commitment (**Fig. 3C**).

## Discussion

During early embryogenesis, MOXD1 is expressed in EMT-transformed and migrating trunk neural crest cells (Fredlund et al., 2023; Knecht and Bronner-Fraser, 2001; Murko et al., 2018). However, the function and possible effects of MOXD1 on these cells at early stages have not been investigated. Here, we show that knockout of *MOXD1* delays development, as quantified by embryo size and somite formation. Whether these differences in growth are retained until hatching remains to be investigated, but our results suggest that MOXD1 is important in early embryogenesis mediated by trunk neural crest cell functions. Further studies should determine the biological pathways and mechanisms that fully govern these effects. Our RNA sequencing data show that *MOXD1* regulates genes involved in cell fate commitment, and imbalances in cell lineage distributions could indeed affect embryogenesis and organ development. In conjunction with this, we have recently shown that *MOXD1* is expressed in a lineage specific pattern during mouse and human SNS development (Fredlund et al., 2023). Bulk- and single cell RNA sequencing demonstrated Schwann cell-restricted expression of *MOXD1*, and in neural crest-derived neuroblastoma, this pattern was reflected by restricted expression in mesenchymal, neural crest- and Schwann cell precursor-like cells (Fredlund et al., 2023).

MOXD1 has all the structural residues and components to bind copper. However, how MOXD1 exert copper-mediated mechanisms, and the impact this might have on embryogenesis, is not known. Our RNA sequencing data showed that copper-related genes *APOA4* and *FAM107A*, the latter a direct binding partner of *COMMD1*, are significantly affected by conditional knockout of *MOXD1* in trunk neural crest cells. We hypothesize that MOXD1 might affect embryo development by skewed copper metabolism in EMT-transformed migrating trunk neural crest cells. Of note, *APOA4* was also affected by *MOXD1* overexpression in neuroblastoma cells devoid of endogenous MOXD1 protein. These results suggest that copper ion binding mechanisms could be important for MOXD1-regulated embryonic development, as well as tumor suppression.

Overlapping genes from RNA sequencing of both healthy neural crest cells and neuroblastoma cells are connected to embryonic development, and extracellular matrix, further strengthening the close relationship between proper control of neural crest and the development of neural crest-derived childhood cancer.

Data from our recent preprint (Fredlund et al., 2023) suggest that MOXD1 is important for prolonged organ development, in particular adrenal glands, tissue derived from trunk neural crest. In conjunction, genes affected by MOXD1 already 24 hours after knockout strongly connect to gland development, suggesting that the role of MOXD1 in regulating proper migration and MOXD1-driven organ formation is pronounced already from the time point of trunk neural crest EMT.

Our combined results demonstrate that MOXD1 is cell lineage specific, plays an important role in neuroblastoma, and is crucial for proper embryonic development. The impact of copper metabolism, and other mechanisms involved, is under present investigation, and will be of importance to understand neural crest development and its derived diseases, including childhood neuroblastoma.

## Material and methods

### Ethics

All chick embryo procedures followed the guidelines set by the Malmö-Lund Ethics Committee for the use of laboratory animals and were conducted in accordance with European Union directive on the subject of animal rights (ethical permit nr. 18743/19).

### Embryos and perturbations

Chick embryos were acquired from commercially purchased fertilized eggs and incubated at 37.5°C until desired developmental Hamburger Hamilton (HH) (Hamburger and Hamilton, 1951) stages were reached. Optimal conditions for high transfection efficiency applying one-sided electroporation *in ovo* were determined to 5 pulses of 30ms each at 22V. Ringer’s balanced salt solution (Solution-1: 144g NaCl, 4.5g CaCl•2H_2_O, 7.4g KCl, ddH_2_O to 500ml; Solution-2: 4.35g Na_2_HPO_4_•7H_2_O, 0.4g KH_2_PO_4_, ddH_2_O to 500ml (adjust final pH to 7.4)) containing 1% penicillin/streptomycin was used in all experiments. All constructs were injected at HH stage 10+/11 into the lumen of the neural tube from the posterior end. Embryos were electroporated *in ovo* in a one-sided fashion. Embryos injected with CRISPR/Cas9 constructs were allowed to sit at room temperature for 6 – 10 hours before further incubation of the embryos at 37.5°C in order to allow the Cas9 protein to fold.

### Constructs

CRISPR constructs with gRNA non-targeting control (#99140, Addgene) or gRNAs targeting *MOXD1* (MOXD1.1.gRNA Top oligo – 5’-cgtcCGGGCCGAACCTATCCGCAC-3’, Bot oligo – 5’-aaacGTGCGGATAGGTTCGGCCCG-3’; MOXD1.2.gRNA Top oligo – 5’-cgtcGGCGTCCGCCGACATCGTCG-3’, Bot oligo – 5’-aaacCGACGATGTCGGCGGACGCC-3’; MOXD1.6.gRNA Top oligo – 5’-cgtcCGGATCTAATGGAGTGCCAA-3’, Bot oligo – 5’-aaacTTGGCACTCCATTAGATCCG-3’) were cloned into U6.3>gRNA.f+e (#99139, Addgene) and electroporated at a concentration of 1.5 ug/ul, and accompanying Cas9-GFP (#99138, Addgene) at 2 ug/ul.

Morpholinos used were from GeneTools with the following sequences, splice modifying MOXD1 oligo (5’-TTTACAGAACATGAGGGTACAAACC-3’) and a corresponding 5’-mispair oligo (5’-TTTAgAcAACATGAcGGaAgAAACC-3’). The MOXD1 morpholino was designed for the exon 6/intron 6 splice boundary. Morpholinos were injected at a concentration of 1mM and co-electroporated with a GFP tagged empty control vector (1 ug/ul).

### Quantification

Embryonic development was determined by counting the number of somites of dissected embryos *ex ovo* or by determining the HH stage of embryos *in ovo* using head and tail morphology. Embryos used for quantification of developmental age were further collected and processed for RNA sequencing. The number of embryos (n) for each group is denoted in the figure legend.

### RNA sequencing

Embryos were incubated at 37.5°C for 36 hours post-electroporation before harvest. The trunk portion of neural tubes were dissected with all other tissue carefully removed, and each neural tube was individually snap frozen. Total RNA was extracted using the RNAqueous Micro Kit (Ambion, #AM1931) according to the manufacturer’s recommendations. In total, samples from 9 control and 12 MOXD1 knockout embryos were collected and analyzed. Sequencing was performed using NovaSeq 6000 system (20012850, Illumina). Alignment of reads was performed using the STAR software (Dobin et al., 2013) and the reference genome was from the Ensemble database (ensemble GRCg6a release 101). Expression counts were performed using the StringTie software. Differentially expressed gene (DEG) analysis was performed using DESeq2 (Love et al., 2014). After removing the not annotated (NA) and LOC-annotated genes, we set the cut-off to p<0.05 and log2foldchange>1.5, ending up with 79 genes. GSEA was performed using clusterProfiler (Wu et al., 2021), with organism set to human, ontology to biological processes, p<0.05, q<0.2 and gene count ≥5.

## Abbreviations

DBH: dopamine beta hydroxylase
EMT: epithelial-to-mesenchymal transition
HH: Hamburger Hamilton
MOXD1: Monooxygenase DBH-like 1
SNS: sympathetic nervous system

## Author contributions

Conceptualization: EF, SM

Methodology: EF, SA, MF, SM

Validation: EF, SA, MF, SM

Formal Analysis: EF, SA, MF, SM

Investigation: EF, SA, MF, SM

Data Curation: EF, SA, MF, SM

Visualization: EF, MF

Funding acquisition: SM

Project administration: SM

Supervision: SM

Writing – original draft: EF, SM

Writing – review & editing: EF, SA, MF, SM

## Data and materials availability

Sequencing data will be made available upon acceptance.

